# Probing and Perturbing Riboswitch Folding Using a Fluorescent Base Analogue

**DOI:** 10.1101/2021.07.22.453416

**Authors:** Janson E. Hoeher, Natalie E. Sande, Julia R. Widom

**Affiliations:** Department of Chemistry and Biochemistry, University of Oregon, Eugene, OR

## Abstract

Riboswitches are mRNA segments that regulate gene expression in response to ligand binding. The Class I preQ_1_ riboswitch consists of a stem-loop and an adenine-rich single-stranded tail (“L3”), which adopt a pseudoknot structure upon binding of the ligand preQ_1_. We inserted 2-aminopurine (2-AP), a fluorescent analogue of adenine (A), into the riboswitch at six different positions within L3. Here, 2-AP functions both as a spectroscopic probe and as a “mutation” that reveals how alteration of specific A residues impacts the riboswitch. Using fluorescence and circular dichroism spectroscopy, we found that 2-AP decreases the affinity of the riboswitch for preQ_1_ at all labeling positions tested, though modified and unmodified variants undergo the same global conformational changes at sufficiently high preQ_1_ concentration. 2-AP substitution is most detrimental to ligand binding at sites proximal to the ligand-binding pocket, while distal labeling sites exhibit the largest impacts on the stability of the L3 domain in the absence of ligand. Insertion of multiple 2-AP residues does not induce significant additional disruptions. Our results show that interactions involving the A residues in L3 play a critical role in ligand recognition by the preQ_1_ riboswitch, and that 2-AP substitution exerts complex and varied impacts on this riboswitch.

## INTRODUCTION

Riboswitches are mRNA segments that regulate gene expression in response to environmental cues, and are most commonly found in bacteria (1). A riboswitch contains an aptamer domain, which binds to a ligand, and an expression platform, which re-folds in response to ligand binding to regulate downstream genes in *cis*. Riboswitches typically control the transcription or translation step of gene expression, although other mechanisms have also been observed (2–4). Since their discovery, riboswitches have been the subjects of intensive study due to their utility as molecular sensors (5, 6) and antibiotic targets (7, 8).

The structures and dynamics of riboswitches are intimately linked to their biological functions. As a result, many techniques have been used to probe riboswitch structure and dynamics, including x-ray crystallography (9), NMR (10, 11), single-molecule Förster resonance energy transfer (smFRET) (12–15) and bulk fluorescence spectroscopy (4, 16). For studies utilizing bulk fluorescence spectroscopy, a powerful approach is to replace a base within the riboswitch with a fluorescent base analogue (FBA), with the most commonly used FBA being the adenine (A) analogue 2-aminopurine (2-AP; Fig. 1) (17). Fluorescence from 2-AP is modulated by its structural context, with stacking on neighboring bases typically inducing quenching (18), and changes in polarity additionally inducing spectral shifts (19). While FBAs are designed to preserve the Watson-Crick base pairing properties of the corresponding native bases, 2-AP has been shown to alter the structure and dynamics even of double-stranded DNA (20). FBAs may impact structural features that rely on non-Watson-Crick interactions to an even greater degree, and such interactions are common in the complex 3-dimensional structures that riboswitches and other RNAs can adopt (21). Some FBAs have been reported that preserve both the Watson-Crick and Hoogsteen faces of adenine, providing effective surrogates in a wide range of structural and enzymatic contexts (22–24), but lack of commercial availability has significantly limited their uptake. Due to this factor and its decades-long record as a useful probe, 2-AP remains by far the most widely used FBA despite having suboptimal structural and photophysical properties.

**Figure 1.**
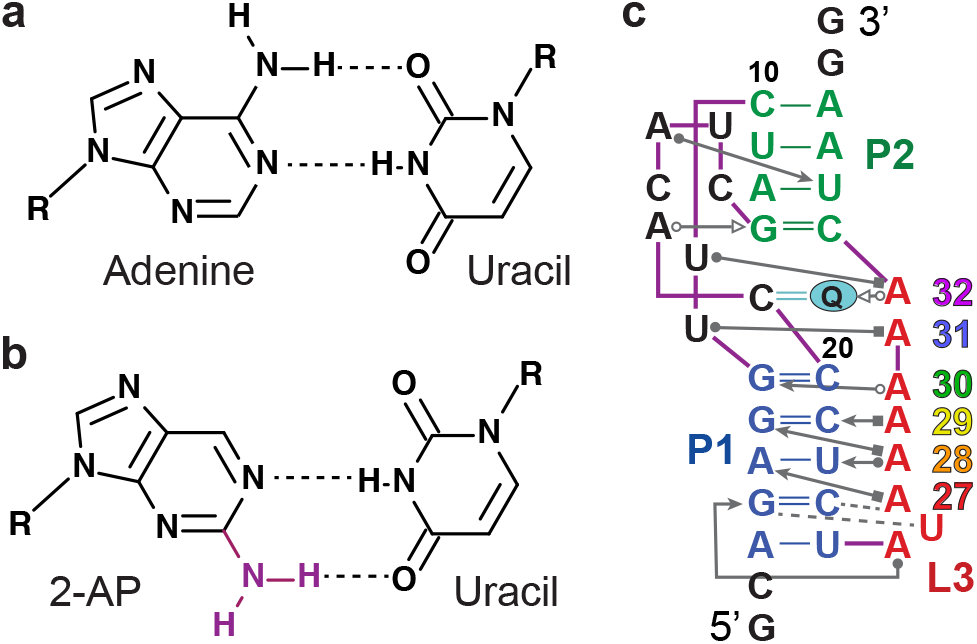
2-Aminopurine and the preQ_1_ riboswitch. (a) Chemical structure of a Watson-Crick base pair between adenine and uracil. (b) Chemical structure of a Watson-Crick base pair between 2-AP and uracil. The re-positioned amine group is shown in purple. (c) The preQ_1_ riboswitch in its docked pseudoknot conformation, shown in Leontis-Westof notation (26). The P1, P2 and L3 domains are indicated in blue, green and red, respectively, and preQ_1_ is shown in cyan. Adenosine residues in L3 are numbered according to the color scheme used in subsequent figures.

To minimize structural perturbations, 2-AP is ideally placed at locations containing non-conserved nucleotides whose hydrogen bonding interactions are not altered by 2-AP substitution. It has been noted that these criteria are satisfied by as many as 10-15% of nucleotides in various riboswitches (25, 26). However, certain sites may be of interest precisely because of their high degree of conservation and non-canonical hydrogen bonding interactions. In these circumstances, 2-AP substitution may act as a perturbing “mutation” that serves concurrently as a spectroscopic probe. If its structural impacts are accounted for, 2-AP can be a useful tool even at highly conserved or structurally complex sites. The approach of using an FBA as a concurrent probe and mutation would be particularly valuable for the study of riboswitches for which high-resolution structures are not available. For such riboswitches, the significance of particular residues is typically inferred through mutagenesis and biochemical methods such as chemical secondary structure probing. FBAs such as 2-AP offer a chemical library of potential “mutations’’ that expands the 3-member library of native base substitutions, and, like the varied reagents used for secondary structure probing (27), different FBAs are sensitive to different aspects of RNA structure (for example, see ref. (28) for a comparison of 2-AP and the cytosine analogue pyrrolocytosine). Combining biochemical and spectroscopic approaches thus offers the potential to expand the level of insight that can be gained in the absence of high-resolution structures.

We utilized 2-AP as a concurrent probe and mutation to study the significance of tertiary interactions involving highly conserved adenosine residues in the Class 1 preQ_1_ riboswitch from *Bacillus subtilis* (*Bsu*; Fig. 1c), which is responsible for regulating genes involved in synthesis of the hypermodified nucleobase queuosine (29). This riboswitch binds to the metabolic intermediate preQ_1_, causing a conformational change that suppresses expression of genes involved in the production of queuosine by enhancing transcription termination (12). Its aptamer domain contains a stem-loop structure (P1) followed by an A-rich tail (L3), and preQ_1_ binding stabilizes a pseudoknot-structured “docked” conformation containing a second helix, P2 (9). The L3 domain of this riboswitch contains a stretch of 6 highly conserved A residues that participate in a complex network of hydrogen bonding with the minor groove of the P1 helix and the ligand-binding pocket (Fig. 1c) (9–11). 2-AP has been utilized extensively in Class I preQ_1_ riboswitches, particularly those from *Thermoanaerobacter tengcongensis* (*Tte*) (30, 31) and *Fusobacterium nucleatum* (*Fnu*) (4), with some prior work in *Bsu* (16), but L3 residues have gone mostly uninvestigated. The kinetics of structural rearrangements of residue A24 in the *Tte* preQ_1_ riboswitch (most closely analogous to A28 in *Bsu*; Fig. 1c) were investigated using 2-AP, revealing almost no change in fluorescence in response to Mg^2+^ and a moderate decrease in response to preQ_1_ (30). However, that base resides in a different sequence context, A(2-AP)C, than the corresponding residue A28 in the *Bsu* riboswitch, A(2-AP)A, and the L3 domain is one nucleotide longer in *Tte*.

We investigated riboswitch variants in which 2-AP replaced subsets of the six consecutive A residues in the L3 domain. Rather than pursuing structurally neutral FBA labeling sites, this work uses 2-AP to introduce site-specific disruptions in the structure. We used fluorescence spectroscopy to investigate how ligand binding affects the local environment at each labeling site, and to determine the affinity of each variant for preQ_1_. These measurements revealed a rich variety of photophysical impacts on 2-AP, which can be attributed to changes in both base stacking and local polarity that occur upon ligand binding (19). Fluorescence lifetime measurements on a subset of variants revealed that 2-AP samples multiple local environments and that ligand binding changes their relative prevalence. We additionally performed absorbance-based measurements that allowed us to directly compare the native riboswitch to the 2-AP-modified variants, which is not possible using fluorescence spectroscopy. Specifically, we used circular dichroism (CD) spectroscopy and thermal denaturation experiments to study the effects of 2-AP and preQ_1_ on the structure and stability of the riboswitch. We found that at every labeling site tested, 2-AP decreases the affinity of the riboswitch for preQ_1_, but the riboswitch still undergoes a global conformational change of a similar nature at sufficiently high ligand concentrations. Placing 2-AP distal to the ligand binding pocket makes the riboswitch more susceptible to thermal denaturation in the absence of preQ_1_, whereas the largest decreases in affinity were observed with 2-AP proximal to the binding pocket. Our results reveal that interactions involving the 6-amino groups of the A residues in L3 are critical for efficient ligand binding by the preQ_1_ riboswitch, and that the formation of these interactions is coupled to changes in base stacking and polarity.

## MATERIALS AND METHODS

RNA oligonucleotides were purchased from Dharmacon (Horizon Discovery) and DNA oligonucleotides were purchased from Integrated DNA Technologies (sequences in Table S1). All modified oligonucleotides were HPLC-purified by the manufacturer, and the 3’ segments were purchased with a 5’ phosphate. Riboswitch samples were prepared by splinted ligation using T4 RNA ligase 2 (New England Biolabs M0239S). 5 nanomoles each of the 5’ and 3’ segments of the riboswitch and the DNA splint were combined and heated to 90 °C for 2 minutes and then allowed to cool to RT over 10 minutes. The manufacturer-provided RNA ligase 2 reaction buffer was added 30 seconds into cooling. After cooling, the mixture was diluted in reaction buffer to a total volume of 150 µL and 60 units of T4 RNA ligase 2 were added. The reactions were incubated at room temperature for 3 hours, then resolved on a 15% polyacrylamide-urea gel. The product band was recovered from the gel by electroelution using a BioRad model 422 electro-eluter followed by ethanol precipitation in the presence of 300 mM NaOAc, pH 5.3. 2-AP riboside was purchased from Tri-Link Biotechnologies.

All spectra were recorded at 20 °C with the sample in a buffer consisting of 20 mM Na_i_PO_4_ at pH 7.5 and 100 mM NaCl. Samples were heated to 90 °C for 2 minutes and then crash-cooled in ice water for 10 minutes before measurement in order to allow the RNA to fold without dimerizing. Where noted, MgCl_2_ was added after cooling to a final concentration of 1 mM. CD spectra and melting curves were recorded on a Jasco J-1500 spectrometer. Melting curves were recorded with the riboswitch at a concentration of 2 µM or AA(2-AP)AC at a concentration of 16 µM. The sample was heated at a rate of 0.75 °C per minute in a 1 cm path-length, low head-space cuvette (Starna 26.160/LHS) while monitoring CD and absorbance at a wavelength of 258 nm. CD spectra were recorded in a 1 cm path-length cuvette (Starna 16.160-10) with the riboswitch at a concentration of 2 µM. Scans were performed from 200 to 350 nm at a rate of 10 nm per minute with an integration time of 4 seconds and an excitation bandwidth of 2 nm, and four sequential scans were averaged together. CD titrations were performed with the riboswitch at a concentration of 50 nM in a 2 cm path-length cuvette. Spectra and melting curves were background-corrected by subtracting off a buffer scan (including preQ_1_ where appropriate). CD spectra were converted from units of millidegrees to extinction coefficient (L/mol*cm) using the following equation, where *CD* is the signal in millidegrees, *C* is the concentration in mol/L and *L* is the path length in cm:

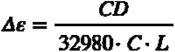

Finally, they were converted to Δε per nucleotide by dividing by the number of nucleotides in the riboswitch, 38.

A fitting procedure was performed in Mathematica (Wolfram Research) to extract melting temperatures (*T_m_*) and transition widths (Δ*T*) from melting curves (33). First, a linear fit was performed to the segment of the melting curve recorded between 75 and 80 °C, and was extrapolated to lower temperature. The absorbance vs. temperature curve was then converted to a proxy for the “fraction folded” at temperature *T*, θ(*T*), using the following equation:

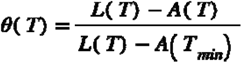

where *L*(*T*) is the value of the linear fit at temperature *T*, *A*(*T*) is the recorded absorbance at temperature *T*, and *A*(*T_min_*) is the recorded absorbance at the lowest temperature. θ(*T*) was then fit using the following equation:

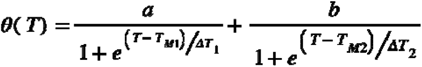

where *a* and *b* are the weights, *T_M1_* and *T_M2_* are the melting temperatures, and Δ*T_1_* and Δ*T_2_* are the widths of unfolding transitions 1 and 2, respectively. Each experimental replicate was fit individually, then the average and standard deviation of the resulting parameter values were determined.

Fluorescence spectra were recorded on an Edinburgh FS5 fluorometer with the RNA at a concentration of 500 nM. After each addition of ligand, the sample was briefly pipette mixed, stirred in the sample holder of the fluorometer for 4 minutes, then allowed to rest for 1 minute before scanning. Scans were performed from 325 to 500 nm with an excitation bandwidth of 5 nm at 305 nm, an emission bandwidth of 2 nm, and an integration time of 1 s. To account for the minor effect of dilution of the RNA due to ligand addition, spectra were corrected according to the following equation:

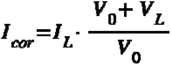

where *I_cor_*is the concentration-corrected intensity, *I_L_* is the uncorrected intensity at ligand concentration *L*, *V_0_* is the initial volume of sample before any ligand was added, and *V_L_* is the cumulative volume of ligand that had been added before *I_L_* was measured.

The absorption spectrum of a 50 µM solution of preQ_1_ was recorded and used to correct fluorescence spectra for the inner filter effect using the following equation:

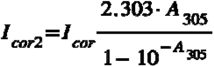

where *I_cor2_* is the concentration- and inner filter effect-corrected intensity and *I_cor_* is the concentration-corrected intensity. *A_305_* is the total absorbance of the sample at the excitation wavelength of 305 nm, calculated according to:

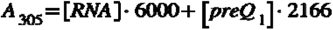

where 6000 L/mol*cm is the extinction coefficient of 2-AP, and 2166 L/mol*cm is the extinction coefficient of preQ_1_ at 305 nm determined from its absorbance spectrum. λ_max_ was extracted by performing a Gaussian fit to the region of the spectrum between 350 and 380 nm.

3 replicates were performed for each fluorescence titration, the corrected fluorescence intensities or λ_max_ values were averaged together, then the resulting curve was converted into fractional saturation (FS; scaled to run between 0 and 1). It was then fit with the following equation to extract an apparent *K_D_*, where *R* is the RNA concentration, *L* is the ligand concentration, and *a* is a scaling factor:

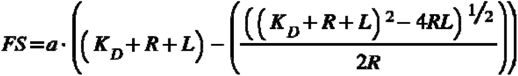

Fluorescence lifetime measurements were performed via time-correlated single photon counting on the Edinburgh FS5 in reverse mode using a 320 nm pulsed LED for excitation. Three replicates were performed for each sample, and a reconvolution fit was performed on each replicate in Edinburgh’s Fluoracle software. A 4-exponential model was required to achieve adequate fits for all riboswitch measurements (Table S2), whereas 2-AP riboside required only a single exponential. Average observed lifetimes ⟨τ⟩_obs_, dark populations α_0_ and corrected amplitudes of decay components α_c_ were computed from the resulting fitting parameters as previously described (18, 34). One of the four exponential components had a lifetime of <100 ps for all samples tested. Its population and lifetime were considered unreliable due to the ~1 ns full width at half maximum of the instrument response function, so it was subsumed into the dark population in downstream analysis.

## RESULTS AND DISCUSSION

The *Bsu* preQ_1_ riboswitch contains six consecutive A residues in its L3 domain, making it a natural system in which to employ 2-AP substitution. We prepared an unmodified riboswitch that lacked 2-AP, singly-labeled variants that contained 2-AP in position 27, 28, 29, 30, 31, *or* 32, as well as doubly-labeled variants containing 2-AP at positions 27 *and* 28, 30 *and* 31, or 31 *and* 32 (Fig. 1c). The bases in L3 are highly conserved (29), suggesting that alteration of them through 2-AP substitution could have detrimental impacts on riboswitch structure and function. Perhaps as a consequence of this, 2-AP labeling sites have been almost entirely limited to other locations in the preQ_1_ riboswitches from both *Bsu* (16) and other organisms (4, 30, 31). To account for this fact, we employed absorbance-based measurements that allowed us to directly compare modified and unmodified riboswitch variants, in addition to fluorescence measurements that exploited the sensitivity of 2-AP to local RNA structure.

### Fluorescence spectroscopy: ligand affinity and comparison of modified variants

2-AP fluorescence exhibited a variety of behaviors at different labeling sites within L3. All riboswitch variants were quenched at least 4-fold compared to free 2-AP riboside, which is unsurprising given that 2-AP fluorescence is quenched as a result of base-stacking (18, 19, 35). Variants in which 2-AP is flanked by two A residues all exhibited similar fluorescence intensities in the absence of Mg^2+^ and preQ_1_, with a 20% difference in intensity between the brightest of them (position 30) and the darkest (position 29) (Fig. 2). Variants with 2-AP flanked by U and A (position 27) or A and C (position 32) both exhibited fluorescence only about half as intense as the other four. A short oligonucleotide containing 2-AP in the same sequence context as A31, AA(2-AP)AC, exhibited a very similar intensity and peak wavelength (λ_max_) to the position 28-31 variants, suggesting that these spectra are characteristic of 2-AP in a generic oligo(A) context rather than being altered significantly by the structure of the riboswitch. This is consistent with previous NMR measurements, which showed that in the absence of divalent cations (like most of our measurements), the sequence of L3 adopts a single-stranded A-form conformation both in the riboswitch and as an isolated 12-mer (36). In contrast, in the presence of Mg^2+^, preorganization of L3 has been observed by MD simulations (37) and smFRET (12, 14). To investigate this further, we recorded spectra of the position 29 and 31 variants in the presence of 1 mM MgCl_2_. While the fluorescence intensity of each variant was nearly unchanged by addition of Mg^2+^, a small (0.7 nm) but resolvable blueshift was observed for position 31 and a larger one for position 29 (2.5 nm), suggesting that 2-AP is in a more nonpolar environment in the presence of Mg^2+^ (19).

**Figure 2.**
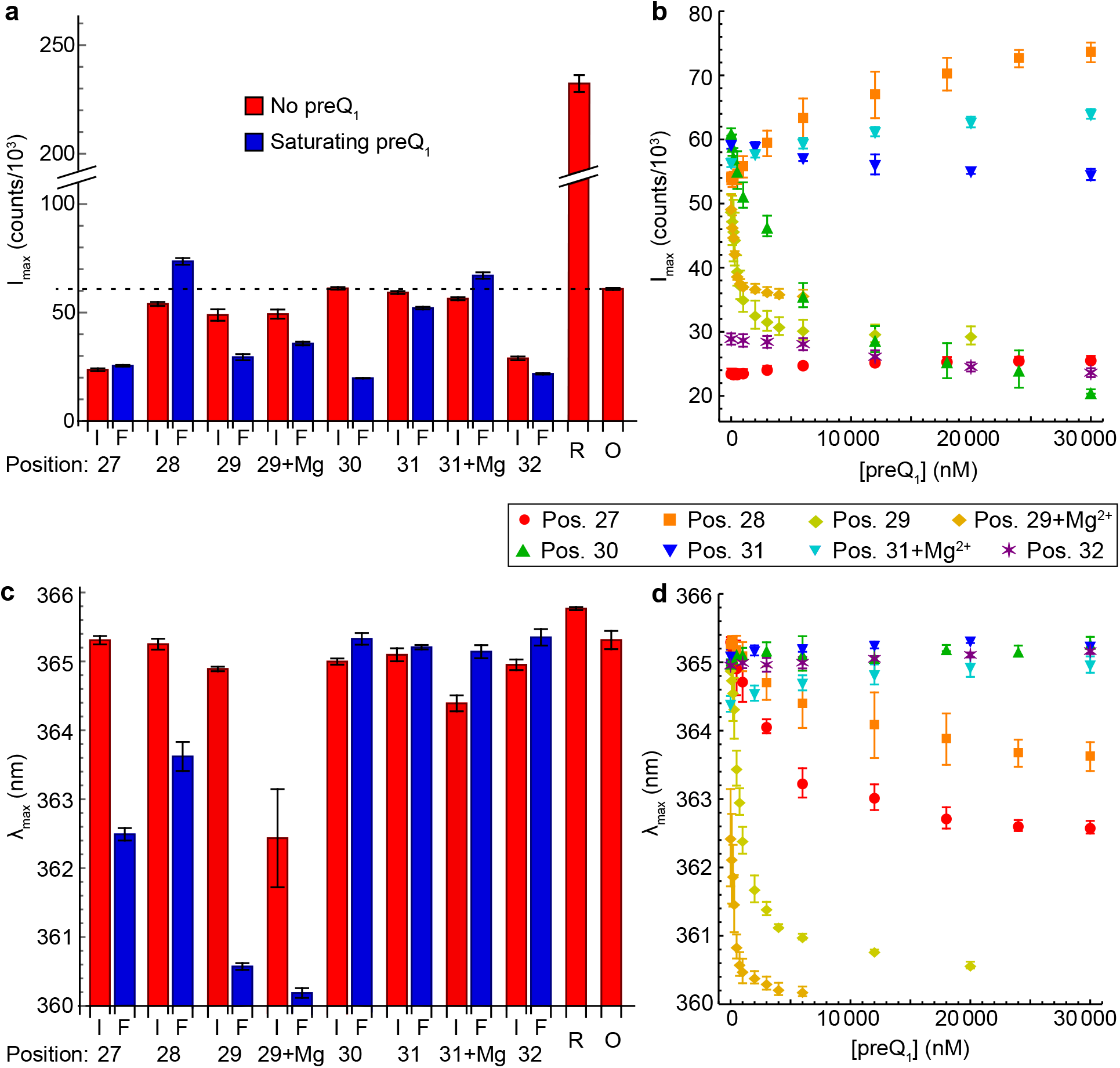
Fluorescence measurements. “+Mg” indicates that the measurement was performed in 1 mM MgCl_2_. (A) Peak fluorescence intensity of each 2-AP-modified riboswitch variant (positions 27-32) at the initial concentration of 0 nM preQ_1_ (I, red) or the final (highest) concentration measured (F, blue). R = 2-AP riboside. O = oligonucleotide AA(2-AP)AC. The dashed line indicates the fluorescence intensity of AA(2-AP)AC. (B) I_max_ vs. [preQ_1_] for each variant over the range of 0 to 30000 nM preQ_1_ (full ranges shown in Fig. S1-S2). (C) Initial and final peak fluorescence wavelengths of riboswitch variants, 2-AP riboside and AA(2-AP)AC. (D) λ_max_ vs. [preQ_1_] for each riboswitch variant. In panels B and D, red: position 27; orange: position 28; yellow: position 29; green: position 30; blue: position 31; purple: position 32. Error bars show the standard deviation across measurements on three samples.

Ligand titrations were performed using fluorescence emission spectroscopy to assess the structural changes induced by preQ_1_ binding and to estimate the affinities of 2-AP-modified variants for preQ_1_ (Figs. 2, S1 and S2, Table 1). A CD titration was performed on the unmodified riboswitch for comparison (Fig. S3). While binding affinities (K_D_) can be obtained more directly by methods that detect thermodynamic changes that occur upon binding, such as microscale thermophoresis (38) and isothermal titration calorimetry (39), 2-AP fluorescence has the benefit of simultaneously reporting on changes in the local environment at the labeling site. In the absence of Mg^2+^, the apparent K_D_ values (K_app_) of 2-AP-labeled variants ranged from 360 nM with 2-AP at position 29 to over 20,000 nM for positions 31 and 32, in stark contrast to the value of 28 nM determined for the unmodified riboswitch via CD. Placing 2-AP at position 28 was quite detrimental (K_app_ = 12,800 nM), which is surprising given that that site is the least conserved of the six (29). The fluorescence intensity and spectrum of each variant responded to preQ_1_ in a distinct manner, though the functional significance of these changes must be considered in light of the dramatic weakening of ligand binding in certain variants. Nevertheless, there was no obvious correlation between the affinity of a variant for preQ_1_ and the magnitude of the changes in its fluorescence intensity or λ_max_, showing that even if 2-AP substitution at a particular site decreases affinity, that residue may still undergo significant structural rearrangements at ligand concentrations sufficiently high for binding. Furthermore, our CD and thermal denaturation measurements (detailed below) indicate that the impacts of preQ_1_ on the global structure and stability of the riboswitch are similar for unmodified and modified variants. These observations suggest that 2-AP fluorescence provides meaningful information about how the local environments of L3 residues differ between the pre-docked and ligand-stabilized docked conformations.

**Table 1.**
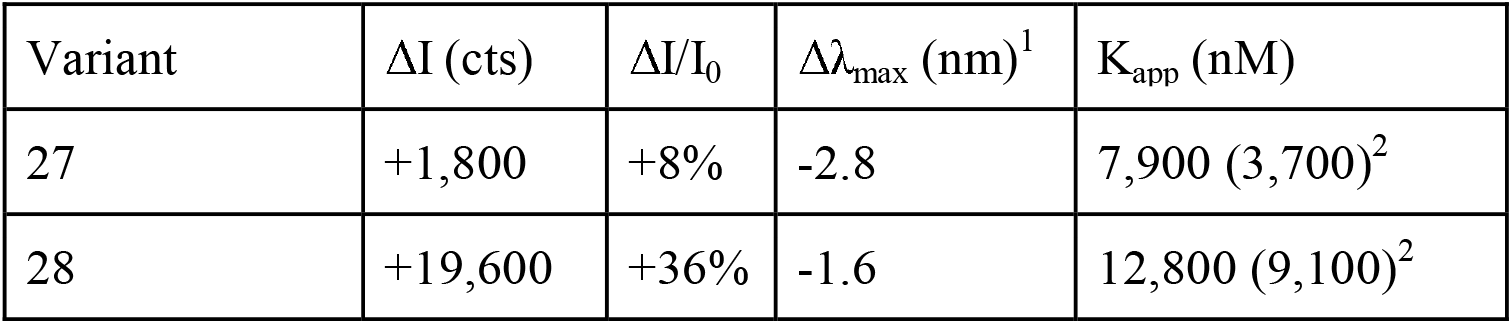
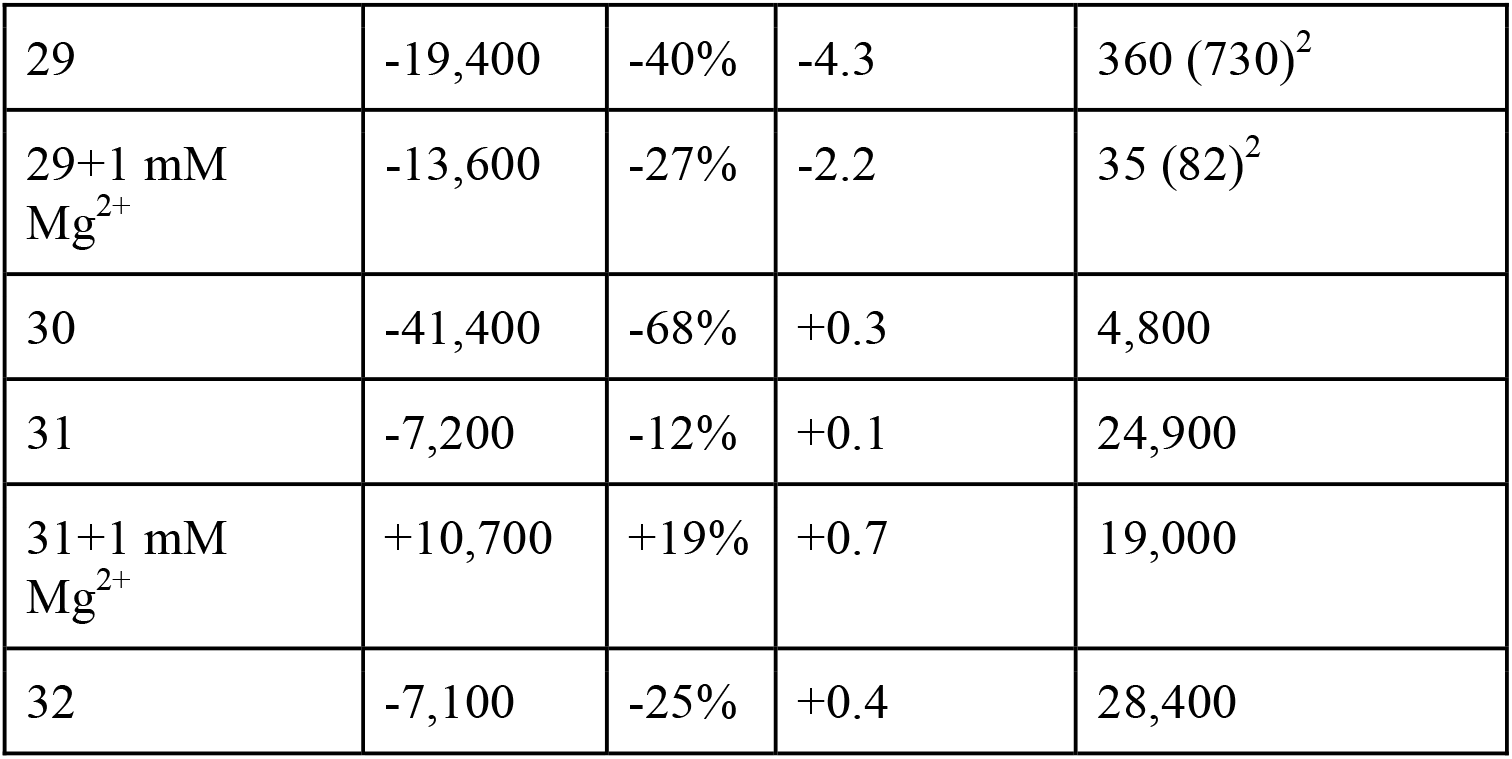
Fluorescence titration results. ΔI: change in maximum intensity from 0 nM preQ_1_ to highest concentration measured. ΔI/I_0_: change in intensity relative to intensity in absence of ligand. Δλ_max_: change in maximum wavelength from starting to ending point. K_app_: apparent K_D_ determined by fitting titration curve. ^1^Typical standard deviations in λ_max_ were ±0.2-0.5 nm. ^2^Based on λ_max_. All other values are based on I_max_.

| Variant | $\Delta I$ (cts) | $\Delta I/I_0$ | $\Delta\lambda_{\max}$ (nm)<sup>1</sup> | $K_{\text{app}}$ (nM) |
| --- | --- | --- | --- | --- |
| 27 | +1,800 | +8% | -2.8 | 7,900 (3,700)<sup>2</sup> |
| 28 | +19,600 | +36% | -1.6 | 12,800 (9,100)<sup>2</sup> |
| 29 | -19,400 | -40% | -4.3 | 360 (730)<sup>2</sup> |
| 29+1 mM<br>Mg<sup>2+</sup> | -13,600 | -27% | -2.2 | 35 (82)<sup>2</sup> |
| 30 | -41,400 | -68% | +0.3 | 4,800 |
| 31 | -7,200 | -12% | +0.1 | 24,900 |
| 31+1 mM<br>Mg<sup>2+</sup> | +10,700 | +19% | +0.7 | 19,000 |
| 32 | -7,100 | -25% | +0.4 | 28,400 |

In addition to having the lowest fluorescence intensities in the absence of preQ_1_, the position 27 and position 32 variants also exhibited the smallest absolute changes in 2-AP fluorescence intensity upon ligand addition, with position 27 exhibiting an increase of only ~1,800 counts (8%). This slight change in intensity was non-monotonic as a function of preQ_1_ concentration (Fig. S4), leading to a titration curve fit that was significantly poorer (R^2^=0.982) than all other variants (R^2^=0.996-0.999). This variant exhibited the second-largest change in λ_max_ upon addition of preQ_1_, and the concentration-dependence of λ_max_ was much better fit (R^2^=0.998) than I_max_. This yielded a K_app_ of 3,700 nM, which is likely to be more reflective of this variant’s ligand binding behavior than the I_max_-based value of 7,900 nM. The most significant changes in fluorescence intensity were observed with 2-AP at positions 28 (+36%), 29 (−40%) and 30 (−68%). Interestingly, the equivalents of our positions 29 and 30 are the only sites in L3 that exhibit reduced scission in the presence of preQ_1_ in in-line probing measurements, indicating an increase in the structural rigidity of the backbone (29). These variants were strongly quenched upon addition of preQ_1_, indicating that this increase in rigidity is accompanied by more extensive stacking on neighboring bases.

The *Tte* preQ_1_ riboswitch was found to tolerate 2-AP substitution at a variety of sites in the presence of 2 mM MgCl_2_ with only modest changes in K_D_, though K_D_ was not determined for any A-to-2-AP substitutions in L3 (30). It is possible that the riboswitch is more tolerant of 2-AP substitution in the presence of Mg^2+^, so we investigated its impact on one strongly binding (position 29) and one weakly binding (position 31) variant. While addition of 1 mM Mg^2+^ to the position 31 variant only modestly improved its K_app_ (from 24,900 to 19,000 nM), it changed the response of 2-AP to preQ_1_ from a slight decrease in intensity and no spectral shift to a slight redshift and increase in intensity. In contrast, position 29 retained the blueshift and quenching that it exhibited in the absence of Mg^2+^, but with a significant improvement of K_app_ from 390 nM to <100 nM.

As noted above, spectral shifts upon ligand addition were observed with 2-AP certain sites. Blueshifts in the emission spectrum occurred upon ligand addition with 2-AP at positions 27 (−2.8 nm), 28 (−1.5 nm) and 29 (−4.3 nm in the absence of Mg^2+^, −2.2 nm in the presence of 1 mM Mg^2+^) (Fig. 2 and Table 1). The other sites showed shifts no greater than 0.7 nm. Position 29 showed the largest blueshift, in keeping with its large change in fluorescence intensity. In stark contrast, position 30, which showed the largest absolute and relative change in intensity, showed no blueshift. It has been shown previously using deoxynucleosides that changes in base-stacking alone do not lead to spectral shifts in 2-AP fluorescence, while a decrease in solvent polarity causes both blueshifts and quenching (19). In the preQ_1_ riboswitch, NMR and x-ray crystal structures show that in the presence of ligand, the bases in L3 stack in sets of 2 (9, 11). A27 and 28 form a parallel stack with one another, as do A29 and 30, as do A31 and 32, with changes in angle existing between each set of stacked bases. A29-32 are nearly co-planar with distal bases, including base pairs in P1 in the case of A29 and A30, and U8 and U9 in the case of A31 and A32 (Fig. 1c). A28 is notably displaced and tilted relative to A29, breaking the chain of stacking and potentially explaining the increase in 2-AP fluorescence upon addition of ligand when it is at position 28. A pattern emerges when these stacking partners are considered as units. In the absence of Mg^2+^, 2-AP fluorescence increases upon addition of ligand in the 27-28 stack, decreases strongly in the 29-30 stack, and decreases slightly in the 31-32 stack. Superimposed on this pattern of fluorescence intensity changes, the three bases distal to the ligand binding pocket (27–29) exhibit blueshifts upon ligand addition while the three proximal bases do not.

The combination of blueshift and de-quenching observed with 2-AP at positions 27 and 28 is surprising, given that increased stacking would appear most likely to accompany a more nonpolar environment, while position 29 exhibits the more intuitive combination of blueshifting and quenching. It is important to note that the preQ_1_ riboswitch is known to sample multiple conformations under the experimental conditions utilized here (12–14), and steady-state emission spectra provide a population-and intensity-weighted average across this ensemble of structures. To investigate the distribution of structures present in solution, we performed fluorescence lifetime measurements using time-correlated single photon counting (TCSPC), focusing on the two variants that showed the largest spectral shifts upon ligand addition, positions 27 and 29 (Table 2 and Fig. S5). Both variants were investigated in the absence and presence of saturating preQ_1_, and the position 29 variant was additionally investigated in the presence of 1 mM MgCl_2_.

**Table 2.** Observed average lifetime (⟨τ⟩_obs_), and populations (α_ic_) and lifetimes (τ_i_) of individual decay components obtained through TCSPC. α_0_ is a dark population whose decay timescale is not resolved (τ < 100 ps). The populations α_ic_ have been corrected to account for the presence of the dark population. Percentages may not add up to 100 due to rounding.

| Sample | Mg<sup>2+</sup> | preQ<sub>1</sub> | $\langle\tau\rangle_{\text{obs}}$ (ns) | $\alpha_0$ (%) | $\alpha_{1c}$ | $\tau_1$ | $\alpha_{2c}$ | $\tau_2$ | $\alpha_{3c}$ | $\tau_3$ |
| --- | --- | --- | --- | --- | --- | --- | --- | --- | --- | --- |
| 2-AP riboside | 0 mM | 0 mM | 8.85 | 0 |  |  |  |  | 100 | 8.85 |
| Position 27 | 0 mM | 0 uM | 3.4 | 74 | 17 | 2.1 | 8 | 5.2 | 1 | 10.4 |
| Position 27 | 0 mM | 42 uM | 2.4 | 60 | 26 | 1.1 | 13 | 4.3 | 1 | 10.8 |
| Position 29 | 0 mM | 0 uM | 3.6 | 49 | 24 | 1.9 | 21 | 4.3 | 5 | 9.0 |
| Position 29 | 0 mM | 20 uM | 2.3 | 50 | 37 | 1.6 | 10 | 3.5 | 2 | 8.9 |
| Position 29 | 1 mM | 0 uM | 3.4 | 45 | 29 | 1.8 | 20 | 4.1 | 6 | 8.9 |
| Position 29 | 1 mM | 20 uM | 2.6 | 49 | 39 | 1.9 | 11 | 4.7 | 1 | 12.8 |

Four exponential components were required to adequately fit the resulting fluorescence decays, which was unsurprising considering the complex decays observed even for systems as small as di- and trinucleotides (18, 40–42). Under all conditions tested, the decay time constants of both variants fell within the ranges of 10.9±2, 4.5±1, and 1.6±0.5 ns, along with a significant dark population with decay timescales too fast to resolve accurately with our instrumentation (Table 2). In all three cases (position 27 without Mg^2+^, and position 29 with and without Mg^2+^), addition of preQ_1_ increased the weight of the 1.6±0.5 ns component (Table 2). The conformation that exhibits this decay may therefore be responsible for the blueshift in fluorescence that occurs upon ligand addition in all three cases. Supporting this interpretation, the increase in weight of the ~1.6 ns component is most significant for the condition that exhibits the largest blueshift: position 29 in the absence of Mg^2+^. It has been previously shown that solvent effects alone are sufficient to shorten the fluorescence lifetime of 2-AP free base from >10 ns in neat H_2_O to ~1.5 ns in neat dioxane (19). The latter value is very similar to the 1.6±0.5 ns lifetime that we attribute to the most nonpolar, “bluest” environment sampled by 2-AP at positions 27 and 29 in the preQ_1_ riboswitch. With 2-AP at position 27, the increase in the ~1.6 ns population occurs through a shift of population away from the dark conformation, explaining this variant’s increase in fluorescence intensity in the presence of preQ_1_. With 2-AP at position 29, it occurs through a shift of population away from the ~4.5 and ~10.9 ns decays in both the presence and absence of Mg^2+^, explaining this variant’s decrease in fluorescence intensity. We conclude that these variants share a conformation in which 2-AP is protected from solvent but only moderately base-stacked, and that their changes in fluorescence intensity are determined by which alternative conformations are depleted upon ligand binding.

### CD spectroscopy and thermal denaturation: comparison of unmodified and modified variants

Absorbance-based measurements provided further insight into the behavior of L3, and into how 2-AP substitution alters the properties of the riboswitch. We recorded CD spectra and melting curves for each variant in the absence and presence of preQ_1_. While it can be challenging to extract quantitative structural information from the ultraviolet CD spectra of structured RNAs, they allowed us to determine whether different variants adopted similar global structures to one another, and whether their global structures changed in a similar manner upon preQ_1_ binding. Melting curves, which were recorded by monitoring CD and absorbance at 258 nm during heating, were used to assess the impacts of 2-AP and preQ_1_ onthe structural stability of the riboswitch. Unfolding is typically indicated by an increase in the absorbance signal and a decrease in the CD signal.

#### CD spectroscopy

We found that in the absence of preQ_1_, the unmodified riboswitch exhibits a CD spectrum that is similar to those reported for other RNAs that adopt a pseudoknot structure (43). When preQ_1_ was added, the CD spectrum exhibited a pronounced shift to more negative values (Fig. 3a), reflecting the shifting of equilibrium from the pre-docked (P2 not intact) to the docked (P2 intact) conformation (12, 13). In the absence of preQ_1_, the spectra of the modified variants are quite similar to one another and slightly less intense than the spectrum of the unmodified riboswitch (Fig. S6a). This difference indicates that 2-AP substitution causes a change in the electronic coupling between nearby bases that gives rise to the CD signal (44, 45). A decrease in the CD intensity upon 2-AP substitution is expected because of the differences in the electronic transitions of A and 2-AP, and should be most apparent when an AAA segment is replaced with an A(2-AP)A segment. Placing 2-AP at position 27 had the smallest impact on the lineshape and intensity of the CD spectrum, consistent with its position in a UAA context rather than an AAA context. These results suggest that all variants adopt similar global structures in the absence of preQ_1_. In the longest-wavelength absorption band of 2-AP (centered at 305 nm), the CD spectra of modified and unmodified variants could not be distinguished from one another, so no attempt was made to extract structural information specific to the 2-AP labeling site.

**Figure 3.**
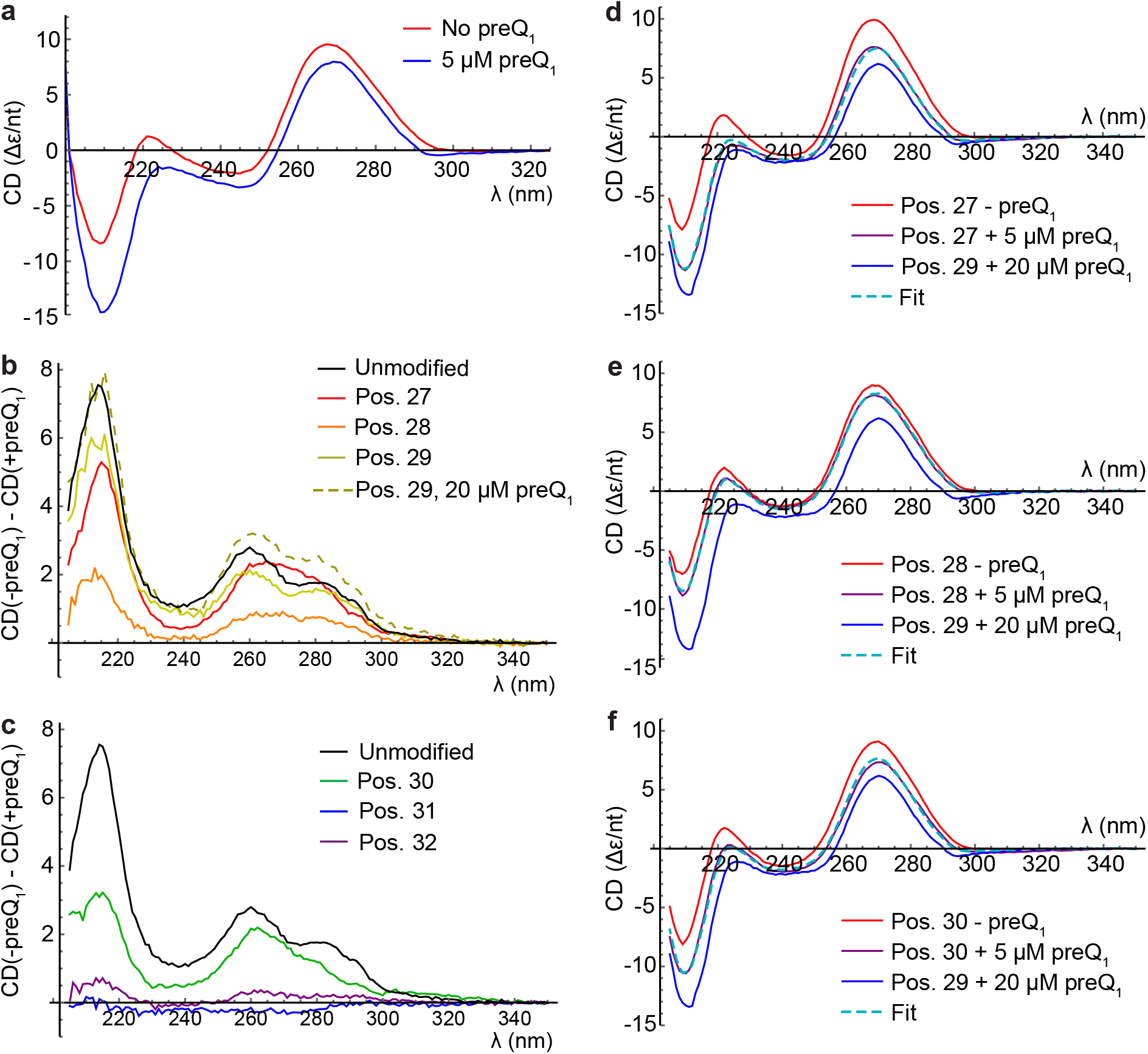
CD analysis of riboswitch variants in the absence of MgCl_2_. (a) Spectra of unmodified riboswitch in the absence (red) or presence (blue) of 5 µM preQ_1_. (b-c) Difference spectra where the CD spectrum recorded in the presence of ligand has been subtracted from the CD spectrum recorded in the absence of ligand. (b) Difference spectra for unmodified riboswitch and variants with 2-AP at positions 27, 28 and 29 using 5 µM preQ_1_, and position 29 using 20 µM preQ_1_. (c) Difference spectra for unmodified riboswitch and variants with 2-AP at positions 30, 31 and 32, all using 5 µM preQ_1_. (d-f) CD spectra fit with linear combinations of −preQ_1_ and Position 29 + 20 µM preQ_1_ spectra. (d) Position 27+ 5 µM preQ_1_ (purple) fit as a linear combination (dashed cyan) of position 27-preQ_1_ (red) and position 29+20 µM preQ_1_ (blue). (e) Corresponding plots for position 28. (f) Corresponding plots for position 30.

Most absorbance-based experiments in the presence of ligand were done with preQ_1_ at a concentration of 5 µM, which based on our K_app_ measurements is saturating for the unmodified riboswitch and nearly saturating for the position 29 variant. The absorption peak of preQ_1_ lies at 260 nm, so for most variants, saturating concentrations of preQ_1_ would have very significant impacts on overall sample absorbance in the UV. For example, for a moderate affinity variant like position 30, a 2 µM RNA sample containing a saturating concentration of 50 µM preQ_1_ would have a total A260 of >1, half coming from preQ_1_. Hence, for most variants, we did not attempt to reach saturation in absorbance-based experiments, but instead considered the trends between the absence of preQ_1_ and partial saturation.

Upon addition of preQ_1_, the spectra of different variants became more distinct, with most exhibiting a general negative shift that was smaller than that exhibited by the unmodified riboswitch (Figs. 3 and S6). The position 29 variant was additionally measured at a saturating preQ_1_ concentration of 20 µM, which induced a shift comparable in magnitude to that exhibited by the unmodified riboswitch (Fig. 3b). If this spectrum is characteristic of a fully preQ_1_-bound riboswitch with a single 2-AP modification, partially saturated riboswitch spectra should be well fit with a linear combination of their respective −preQ_1_ spectra and the saturated position 29 spectrum. This is indeed what was found for the variants with moderate responses to 5 µM ligand (positions 27, 28 and 30; Fig. 3d-f). The fits predict that positions 27, 28 and 30 are 63%, 24% and 49% bound, respectively, in the presence of 5 µM preQ_1_, consistent with the trend in their K_app_ values (Table 1). Even the weakest binder based on K_app_, the position 32 variant, exhibited negative shifts in the CD upon ligand addition at the same wavelengths as the unmodified riboswitch (Fig. 3c). This suggests that much of the variation in +preQ_1_ CD spectra is due to incomplete saturation, and that these variants are capable of undergoing conformational changes of a similar nature to the unmodified riboswitch at sufficiently high ligand concentrations. 2-AP modification at positions 27-30 and 32 thus alters the affinity of the riboswitch for preQ_1_ without altering the fundamental global structures it is capable of adopting.

In contrast, the CD spectrum of the second-weakest binder (position 31) exhibited a small *positive* shift upon ligand addition (Fig. 3c), suggesting that this variant does not undergo the same global conformational change as the others. We investigated this further by recording CD spectra and fluorescence titrations on this variant in the presence of 1 mM MgCl_2_, which was expected to counteract the disruptive 2-AP substitution. Indeed, in the presence of 1 mM Mg^2+^, preQ_1_ addition induced a small negative shift in the CD spectrum at the same wavelengths as observed in the absence of Mg^2+^ for all other variants (Fig. 4c). The fluorescence data reported above show that restoration of this “native’’ global conformational change significantly alters how the local environment of 2-AP at position 31 changes upon ligand binding, while only modestly improving the K_app_ value (from 24,900 to 19,000 nM). This indicates that while 1 mM Mg^2+^ alters the global (based on CD) and local (based on fluorescence) conformations adopted by this variant, it is unable to compensate for the decrease in affinity induced by 2-AP substitution. In contrast, the position 29 variant maintains the same pattern of fluorescence quenching, blueshift in emission spectrum, and negative shift in CD spectrum (Fig. 4b) upon ligand addition regardless of whether 1 mM Mg^2+^ is present, but with a significant improvement in K_app_ in the presence of Mg^2+^.

**Figure 4.**
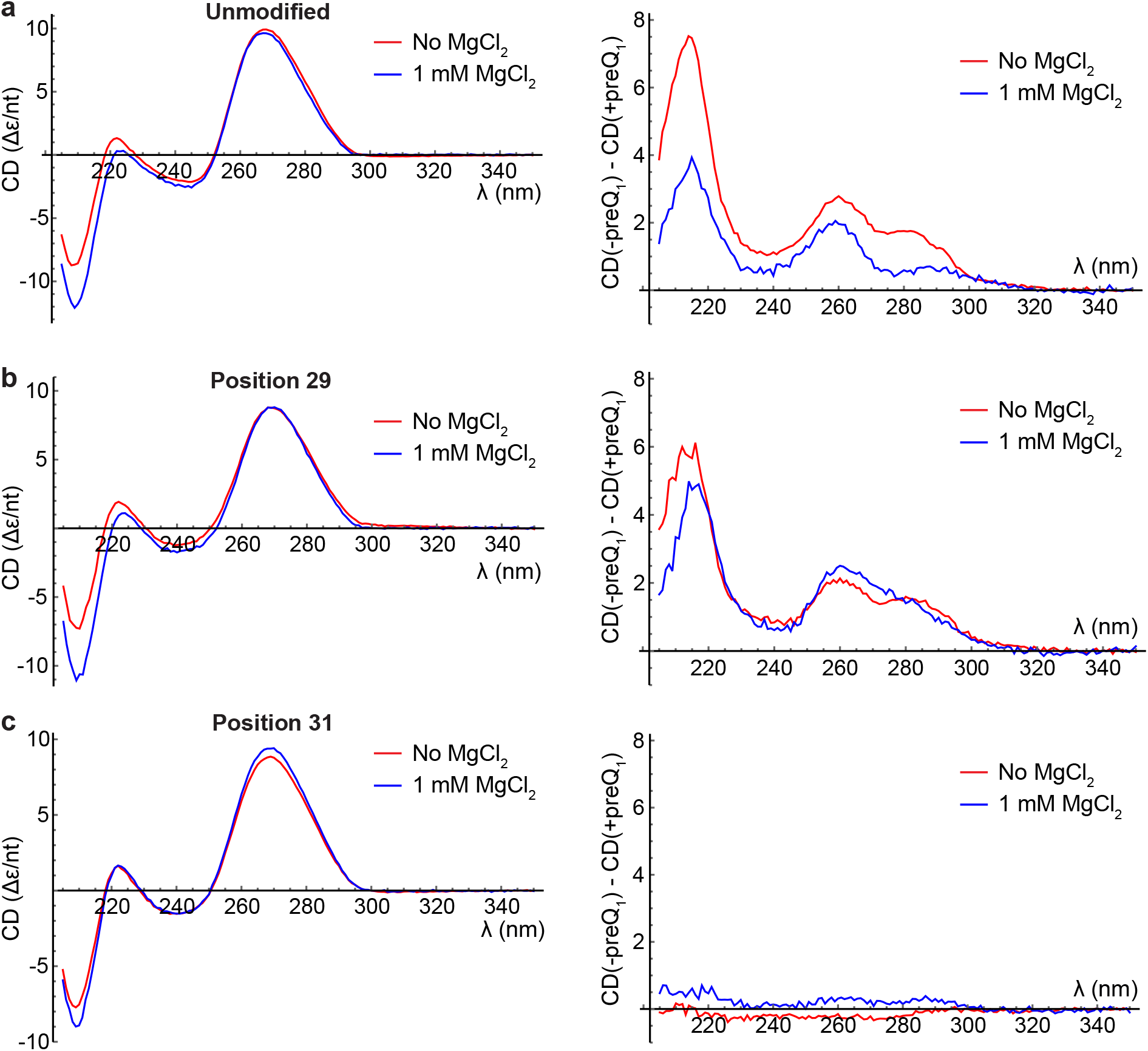
Effects of MgCl_2_ on unmodified, position 29 and position 31 variants. Red curves are reproduced from Fig. 3 for ease of comparison. (a) Left: CD spectra of unmodified riboswitch in the absence of preQ_1_ and the absence (red) or presence (blue) of 1 mM MgCl_2_. Right: CD difference spectra using 5 µM preQ_1_ for unmodified riboswitch in the absence (red) or presence (blue) of 1 mM MgCl_2_. (b) Analogous plots for position 29 variant. (c) Analogous plots for position 31 variant.

CD and fluorescence both indicate that 2-AP substitution at positions 31 and 32 is particularly detrimental, while substitution at position 28 is surprisingly detrimental given its distance from the ligand-binding pocket. NMR (Fig. 5) and x-ray crystallography structures indicate that A31 normally forms a Hoogsteen base pair with U8, which in turn makes a hydrogen bond with preQ_1_ (9–11). Relocation of the 6-amino group of A31 through 2-AP-substitution removes one of the two hydrogen bonds in this Hoogsteen base pair, destabilizing this network of interactions (Fig. 5b). Relocation of the 6-amino group of A32 is expected to break a hydrogen bond with preQ_1_ (Fig. 5c). In the crystal structure, A32 additionally makes a Hoogsteen base pair with U9 (9), but this interaction is not observed in the NMR structure which, like our data on this variant, was recorded in the absence of divalent cations (11)

**Figure 5.**
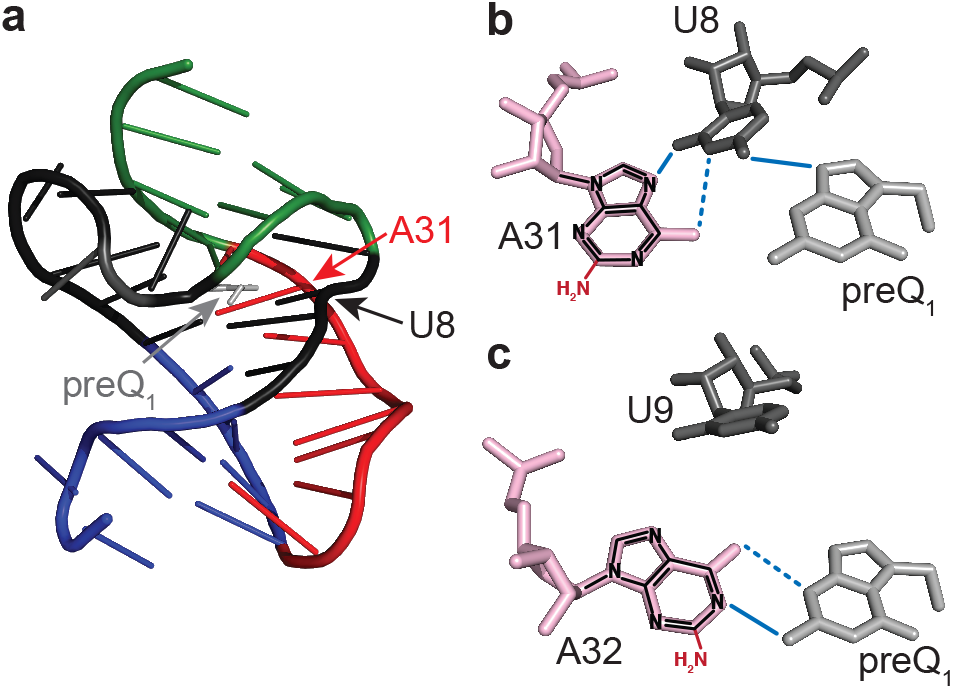
2-AP disrupts key hydrogen bonds when placed at position 31 or 32. (a) NMR structure of the riboswitch (PDB ID 2L1V) with the locations of preQ_1_, U8 and A31 indicated. (b) Closeup of the ligand-binding pocket showing interactions between U8, A31 and preQ_1_. Hydrogen bonds are indicated by blue lines, with dashed lines indicating bonds that are broken when A is replaced by 2-AP. (c) Closeup of the ligand binding pocket viewed from the same perspective showing interactions between A32 and preQ_1_.

Among the labeling sites that are not immediately adjacent to the ligand-binding pocket (positions 27-30), placing 2-AP at position 28 is particularly detrimental to ligand recognition. NMR and crystal structures indicate that each A residue within L3 makes tertiary interactions with residues in P1 (9–11). However, only in A28 does the 6-amino group make hydrogen bonds with members of two different base pairs in P1: U22, which base-pairs with A5, and G6, which base-pairs with C21. Replacing A28 with 2-AP would break these bridging hydrogen bonds, whereas replacing A27, A29 or A30 with 2-AP breaks hydrogen bonds that do not bridge multiple base pairs in P1. This may explain why substitution at position 28 is more detrimental to the affinity of the riboswitch for preQ_1_ than might be expected given its position.

#### Thermal denaturation experiments

To assess the effects of 2-AP substitution on the stability of the riboswitch structure, thermal denaturation measurements were performed by monitoring CD and absorbance signals at 258 nm as the sample was heated. The melting curves of the riboswitch exhibit two transitions (Fig. 6). Based on its breadth and low temperature of onset, we attribute the first to the helix-to-coil transition within L3 (as observed in ref. (36) for a model oligonucleotide mimicking L3), and the higher-temperature transition to unfolding of the P1 helix. A mutant in which the P2 helix is destabilized, G13U, still exhibits two transitions, indicating that transition 1 is not related to unfolding of P2 (Fig. 6a). Furthermore, the oligonucleotide AA(2-AP)AC (described previously in the context of fluorescence measurements) shows a non-cooperative increase in A258 at low temperatures with a steeper slope than the unmodified or mutant riboswitches, but with a shallower slope than certain 2-AP-modified variants (Fig. 6b). This suggests that 2-AP substitution at certain sites destabilizes the single-stranded helical structure of L3.

**Figure 6.**
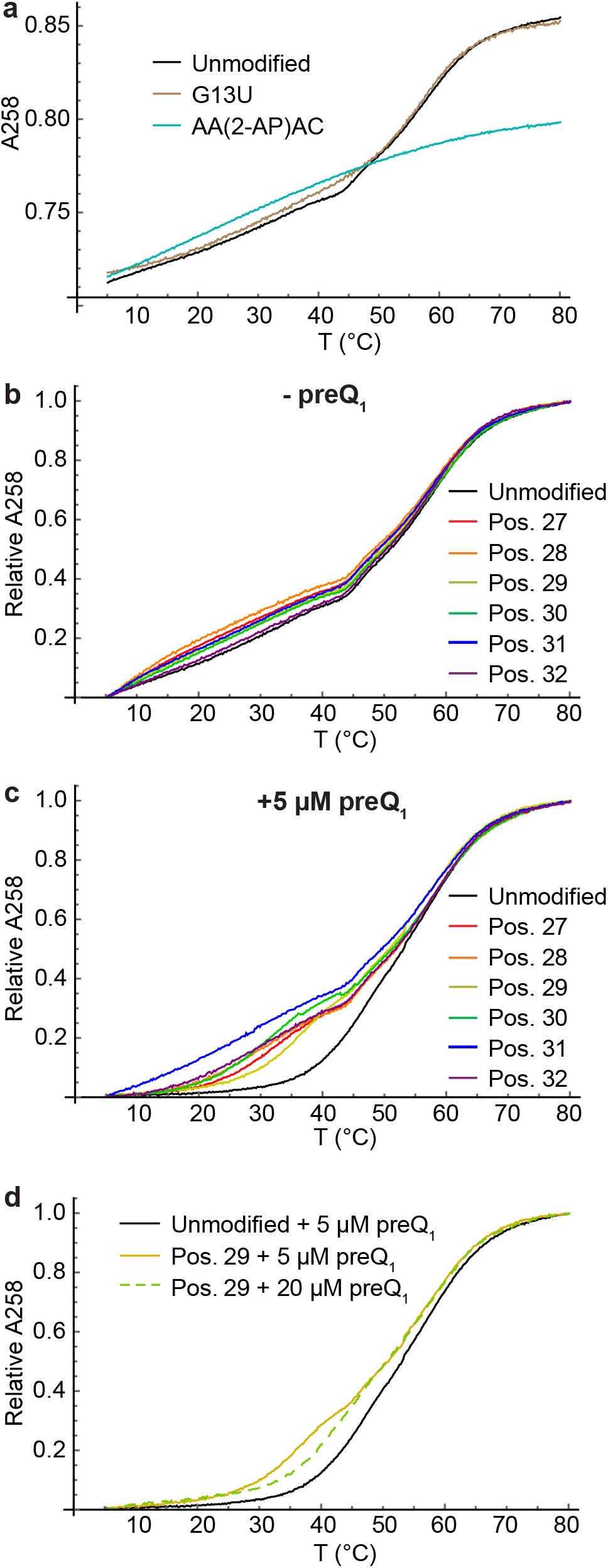
Thermal denaturation measurements. (a) Raw absorbance melting curves of unmodified riboswitch (black) and controls (G13U mutant in brown, AA(2-AP)AC in cyan). (b-c) Absorbance melting curves of unmodified and 2-AP-modified variants recorded at 258 nm in the absence (b) or presence (c) of 5 µM preQ_1_. For ease of comparison, the curves were shifted and normalized to run from a minimum of 0 to a maximum of 1. (d) Rescaled melting curves of unmodified riboswitch (black) and position 29 variant (yellow solid) in the presence of 5 µM preQ_1_, and position 29 variant in the presence of 20 µM preQ_1_ (yellow dashed).

In the absence of preQ_1_, clear differences between the variants are noticeable at low temperatures and apparent when the curves are fit (Fig. 7 and Fig. S7). Transition 1 occurs at T_m_ = 24 ± 2°C in the unmodified riboswitch and at a lower temperature in the 2-AP-modified variants, having a value of T_m_ = 10.2 ± 0.1°C with 2-AP at position 27. T_m_ 1 increases gradually as 2-AP is moved toward the ligand binding pocket to 22.3 ± 0.4°C with 2-AP at position 32. This indicates 2-AP disrupts L3 most significantly when it is placed near the “hinge” where the two domains adjoin, and less significantly when it is near the ligand-binding pocket. 2-AP has little effect on T_m_ 2 (Fig. 7c), which is expected because L3 is already in a random coil conformation at that temperature.

**Figure 7.**
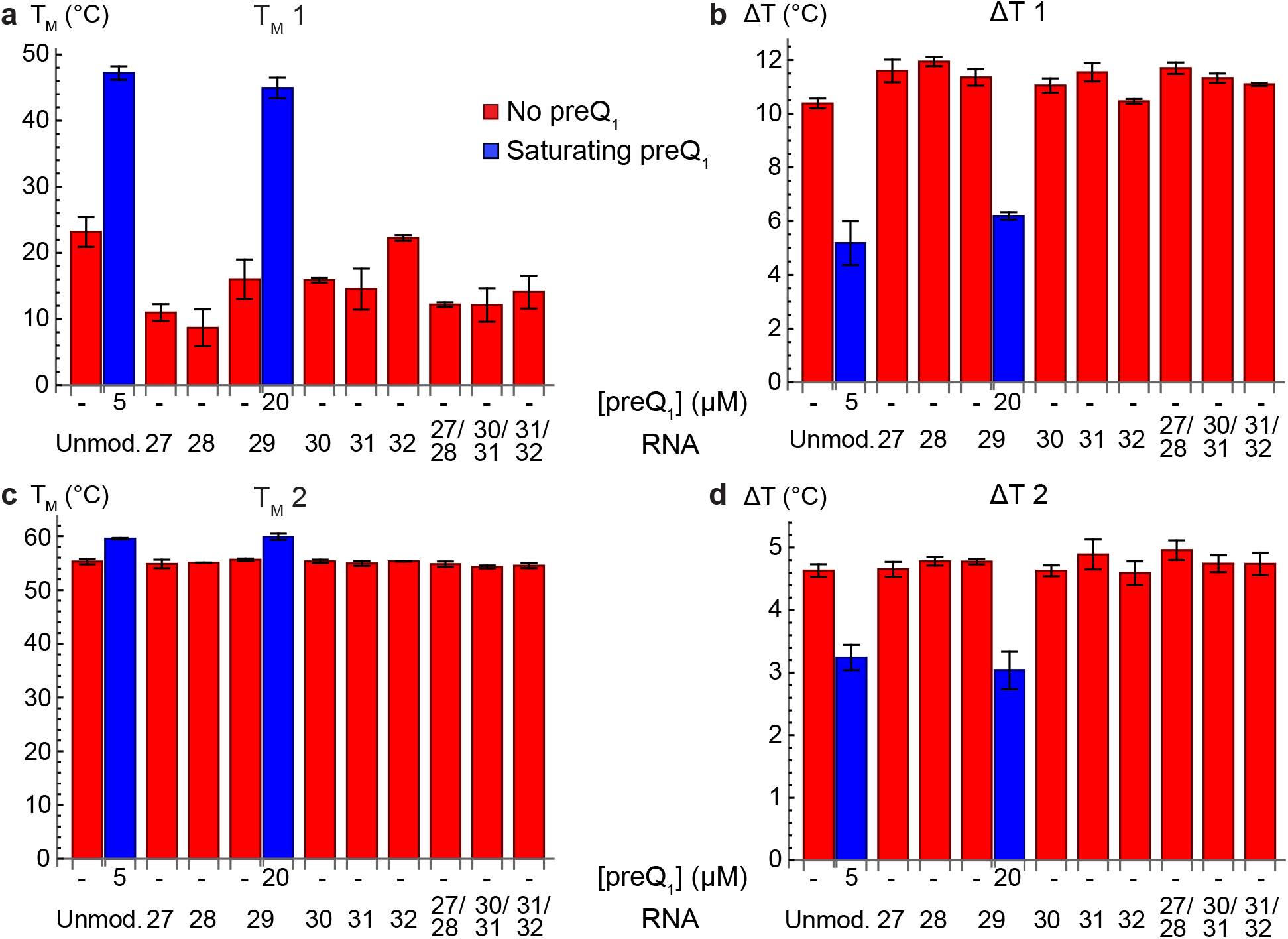
Fitting parameters extracted from absorbance melting curves. Three experimental replicates were fit individually, and the average and standard deviation of each parameter value are plotted. (a) Melting temperature of transition 1 for the unmodified riboswitch (“Unmod.”), singly-labeled variants (27-32), and doubly-labeled variants (27/28, 30/31, and 31/32). Values in the presence of saturating preQ_1_ are shown for unmodified (5 µM) and position 29 (20 µM) variants. (b) Width of transition 1. (c) Melting temperature of transition 2. (d) Width of transition 2.

When preQ_1_ is added to the unmodified riboswitch, transition 1 shifts to a significantly higher temperature, from T_m_ = 24 ± 2°C to 48 ± 1°C; Fig. 6c and Fig. 7) and narrows from ΔT = 10.4 ± 0.2°C to 5.6 ± 0.6°C. This suggests that the L3-P1 interactions that exist in the ligand-bound structure suppress the low-temperature helix-to-coil transition. The melting temperature of transition 2 changes only slightly (from 55.4 ± 0.6°C to 59 ± 1.2°C, while its width decreases to a smaller extent than seen for transition 1 (from 4.7 ± 0.1°C to 3.5 ± 0.1°C; Fig. 6c-d). Addition of saturating (20 µM) preQ_1_ to the position 29 variant induces nearly identical changes in the fitting parameters, though transition 1 remains very slightly lower and broader than in the unmodified riboswitch (Fig. 6d). All other variants exhibit intermediate effects of preQ_1_ that are consistent in magnitude with the effects seen by CD, with 5 µM preQ_1_ inducing almost no change in the melting curves of the position 31 and 32 variants and moderate changes in the melting curves of the position 27, 28 and 30 variants. As with the CD spectra, this is due at least in part to the fact that these variants are not saturated at 5 µM preQ_1_.

When the CD signal is monitored at 258 nm in the absence of preQ_1_, all variants exhibit a monotonic decrease in intensity as the temperature is increased, as expected. In the presence of preQ_1_, all variants except for position 31 exhibit an increase in CD intensity, followed by a decrease starting around 45 °C (Fig. S8). To investigate this surprising observation, we recorded CD spectra at different temperatures during thermal ramping. We observed that in the presence of preQ_1_, the lineshape of the CD spectrum changes with temperature in a manner that leads to a transient increase in CD intensity at 258 nm, while the overall intensity of the spectrum decreases monotonically. As a result, the CD signal at 258 nm is not a simple reflection of the fraction of RNA still folded. Above a temperature of 40 °C, the spectra recorded in the absence and presence of preQ_1_ no longer differ from one another, suggesting that ligand dissociation happens around this temperature.

#### Multiple 2-AP labeling sites

We next investigated whether replacement of multiple A residues with 2-AP would have an additive or cooperative effect on riboswitch folding. We recorded CD spectra and melting curves on variants with 2-AP placed at positions 27 *and* 28, or positions 30 *and* 31, or positions 31 *and* 32 (Fig. S9). In general, a second 2-AP substitution had little impact beyond the more influential of the two single substitutions. For example, the CD spectrum of the position 27 variant exhibits a strong response to 5 µM preQ_1_, while the position 28 variant exhibits a moderate response. When 2-AP is placed at both positions, the response is nearly identical to what is observed in the position 28 variant (Fig. S9a). The same observation is noted when comparing melting curves of singly- and doubly-labeled variants. In the absence of preQ_1_, the absorbance melting curve of each doubly-labeled variant is indistinguishable by eye from that of the more influential singly-labeled variant (Fig. S9d-f), and fitting of the melting curves supports this observation (Fig. 6). A slight additional impact of the second 2-AP residue is observed in melting curves recorded in the presence of 5 µM preQ_1_ (Fig. S9g-i).

### Conclusions

The work presented here comprises a systematic study of the structural changes in the A-rich tail of the preQ_1_ riboswitch that occur upon ligand binding, and how chemical modifications within this domain impact the structure and stability of the riboswitch. The significant impairment of ligand affinity by 2-AP substitution shows that strong ligand binding is dependent on interactions involving the 6-amino group of every A residue in L3. While L3 behaves essentially as a single-stranded RNA in the absence of preQ_1_, ligand binding redistributes the A residues within it between local environments with variable base-stacking and polarity. This work shows that the interactions between L3 and P1 observed in structural studies are critical to high-affinity ligand binding, and that valuable information can be obtained by using FBAs to simultaneously probe and perturb RNA structure.

## ACKNOWLEDGMENTS

This work has been supported in part by NIH R00 GM120457.

## SUPPLEMENTARY MATERIALS

Tables S1-S2 and Figures S1-S9 can be found at DOI: 10.1562/2006-xxxxxx.s1.

