## Supplemental tables 1-2 and figures 1-9 for "Probing and Perturbing Riboswitch Folding Using a Fluorescent Base Analogue"

**Contents**

**Table S1.** Sequences of oligonucleotides

**Table S2.** Goodness of fit for 3- and 4-exponential TCSPC models

**Figure S1.** Fluorescence titration spectra and fits in the absence of $MgCl_2$

**Figure S2.** Fluorescence titration spectra and fits in the presence of $MgCl_2$

**Figure S3.** CD titration on unmodified riboswitch

**Figure S4.** Fluorescence titrations based on peak wavelength

**Figure S5.** TCSPC measurements

**Figure S6.** Raw CD spectra of all variants

**Figure S7.** Melting curve fits

**Figure S8.** CD melting curves and temperature-dependent spectra

**Figure S9.** CD spectra and melting curves of doubly-labeled variants

| | |
|---|---|
| Unmodified | 5’ GCA GAG GUU CUA GCU ACA \| CCC UCU AUA AAA AAC UAA GG |
| Position 27 | 5’ GCA GAG GUU CUA GCU ACA \| CCC UCU AU**(2-AP)** AAA AAC UAA GG |
| Position 28 | 5’ GCA GAG GUU CUA GCU ACA \| CCC UCU AUA **(2-AP)**AA AAC UAA GG |
| Position 29 | 5’ GCA GAG GUU CUA GCU ACA \| CCC UCU AUA A**(2-AP)**A AAC UAA GG |
| Position 30 | 5’ GCA GAG GUU CUA GCU ACA \| CCC UCU AUA AA**(2-AP)** AAC UAA GG |
| Position 31 | 5’ GCA GAG GUU CUA GCU ACA \| CCC UCU AUA AAA **(2-AP)**AC UAA GG |
| Position 32 | 5’ GCA GAG GUU CUA GCU ACA \| CCC UCU AUA AAA A**(2-AP)**C UAA GG |
| Position 27/28 | 5’ GCA GAG GUU CUA GCU ACA \| CCC UCU AU**(2-AP) (2-AP)**AA AAC UAA GG |
| Position 30/31 | 5’ GCA GAG GUU CUA GCU ACA \| CCC UCU AUA AA**(2-AP) (2-AP)**AC UAA GG |
| Position 31/32 | 5’ GCA GAG GUU CUA GCU ACA \| CCC UCU AUA AAA **(2-AP)(2-AP)**C UAA GG |
| G13U | 5’ GCA GAG GUU CUA **U**CU ACA \| CCC UCU AUA AAA AAC UAA GG |
| Control oligo | 5’ AA**(2-AP)**AC |
| DNA splint | 5’ GTT TTT TAT AGA GGG TGT AGC TAG AAC C |

**Table S1.** Sequences of oligonucleotides used in this work. Bold text highlights the positions of 2-AP substitutions or mutations. Vertical lines indicate the division between the 5’ and 3’ segments that were ligated together. Each 3’ segment was purchased with a phosphate group at its 5’ end to enable ligation.

| Sample | $Mg^{2+}$ | $preQ_1$ | $\langle\chi^2\rangle$ (3-exp) | $\langle\chi^2\rangle$ (4-exp) |
|---|---|---|---|---|
| Position 27 | 0 mM | 0 µM | 1.28 | 1.22 |
| Position 27 | 0 mM | 42 µM | 1.37 | 1.20 |
| Position 29 | 0 mM | 0 µM | 1.28 | 1.17 |
| Position 29 | 0 mM | 20 µM | 1.48 | 1.27 |
| Position 29 | 1 mM | 0 µM | 1.34 | 1.22 |
| Position 29 | 1 mM | 20 µM | 1.31 | 1.20 |

**Table S2.** Reduced $\chi^2$ values obtained from 3- and 4-exponential reconvolution fits for each TCSPC dataset. Each value is the average across three replicates that were fit individually.

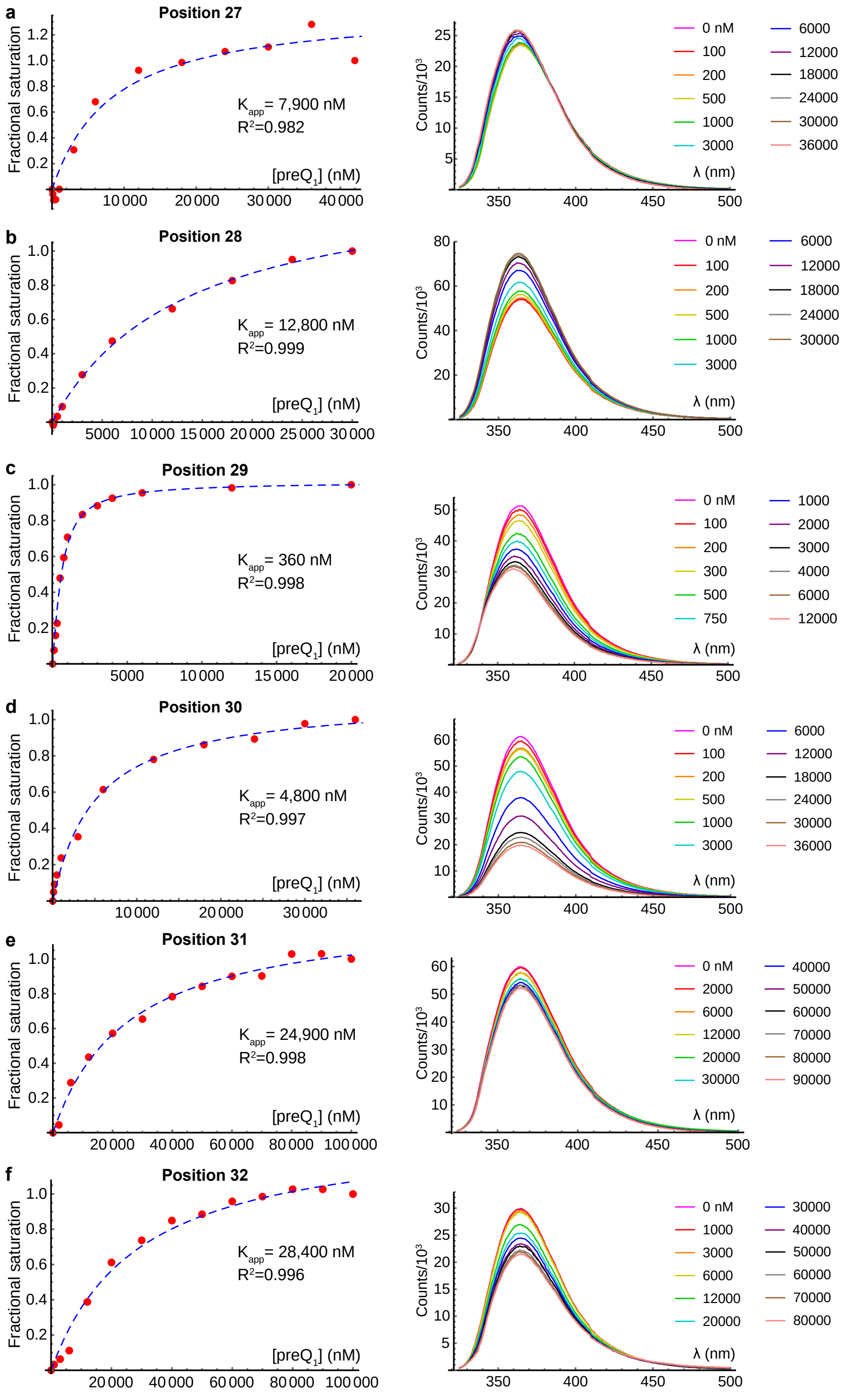


**Figure S1.** Fluorescence titrations on 2-AP-modified variants recorded in the absence of $MgCl_2$. (a) Position 27. (b) Position 28. (c) Position 29. (d) Position 30. (e) Position 31. (f) Position 32. Left: fractional saturation based on peak intensity (red markers), and fit (dashed blue line). The apparent $K_D$ and $R^2$ for the fit are indicated in each plot. Fractional saturation datapoints were computed using the average of three replicates performed on separately prepared samples. Right: Example titration for each variant. A maximum of 12 spectra are shown.

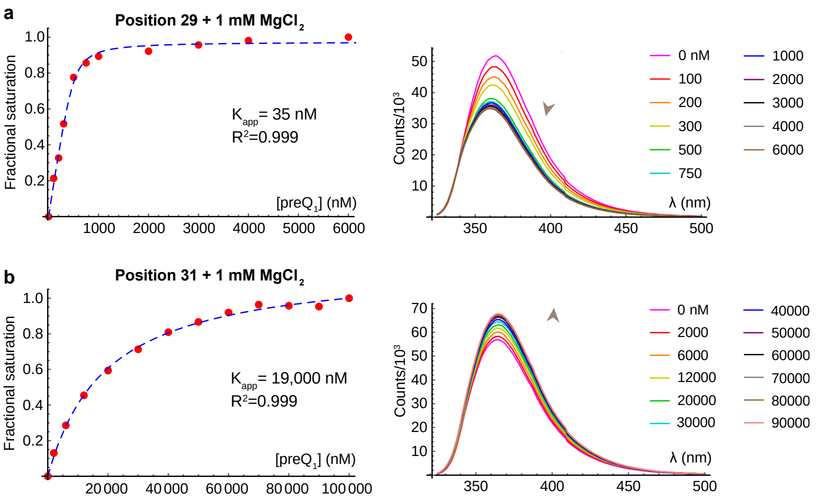


**Figure S2.** Fluorescence titrations on select 2-AP-modified variants recorded in the presence of 1 mM $MgCl_2$. (a) Position 29. (b) Position 31. Left: fractional saturation based on peak intensity (red markers), and fit (dashed blue line). The apparent $K_D$ and $R^2$ for the fit are indicated in each plot. Fractional saturation datapoints were computed using the average of three replicates performed on separately prepared samples. Right: Example titration for each variant. A maximum of 12 spectra are shown.

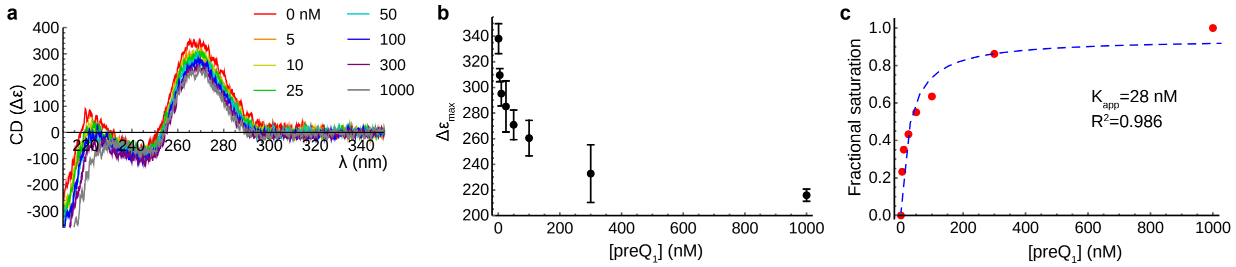


**Figure S3.** CD titration on unmodified riboswitch. (a) CD spectra of 50 nM unmodified riboswitch at varying $preQ_1$ concentrations. (b) Quantification of maximum CD signal at each $preQ_1$ concentration. The average and standard deviation of measurements on three separately prepared samples are shown. (c) Fractional saturation (red markers) and fit (blue dashed line) determined from the data in panel b.

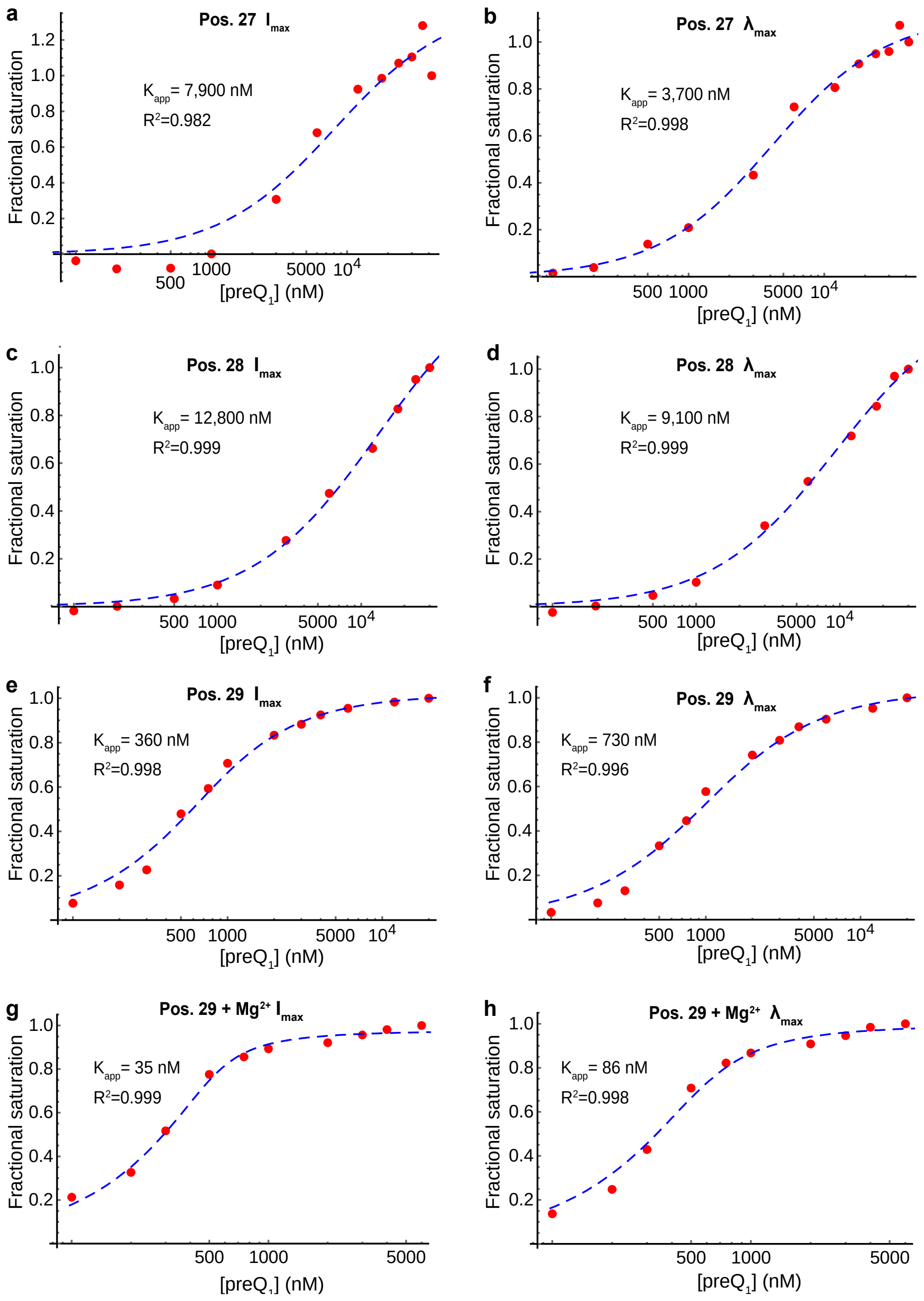


**Figure S4.** Comparison of fluorescence titrations based on peak intensity and peak wavelength. All plots have the x-axis on a log scale to better visualize low concentration datapoints. (a-b) Fractional saturation vs. [preQ$_1$] (red markers) based on peak intensity (a) or peak wavelength (b), and associated fits (blue dashed lines) for position 27 variant. Apparent $K_D$ and $R^2$ for the fits are shown. Note the drastically improved fit in panel b. (c-d) Fractional saturation vs. [preQ$_1$] based on peak intensity (c) or peak wavelength (d) and associated fits for position 28. (e-f) Fractional saturation vs. [preQ$_1$] based on peak intensity (e) or peak wavelength (f) and associated fits for position 29. (g-h) Fractional saturation vs. [preQ$_1$] based on peak intensity (g) or peak wavelength (h) and associated fits for position 29 in the presence of 1 mM $MgCl_2$. All other data was recorded in the absence of $MgCl_2$.

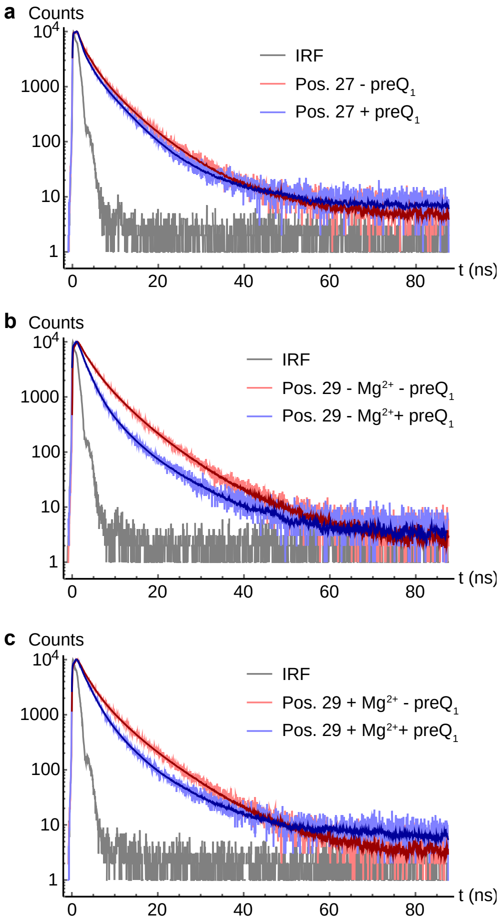


**Figure S5.** TCSPC measurements. (a) Position 27 variant with and without 42 µM $preQ_1$. (b) Position 29 variant with and without 20 µM $preQ_1$. (c) Position 29 variant with and without 20 µM $preQ_1$ in the presence of 1 mM $MgCl_2$. In all panels, gray: instrument response function; red: no $preQ_1$ data (light) and fit (dark); blue: saturating $preQ_1$ data (light) and fit (dark).

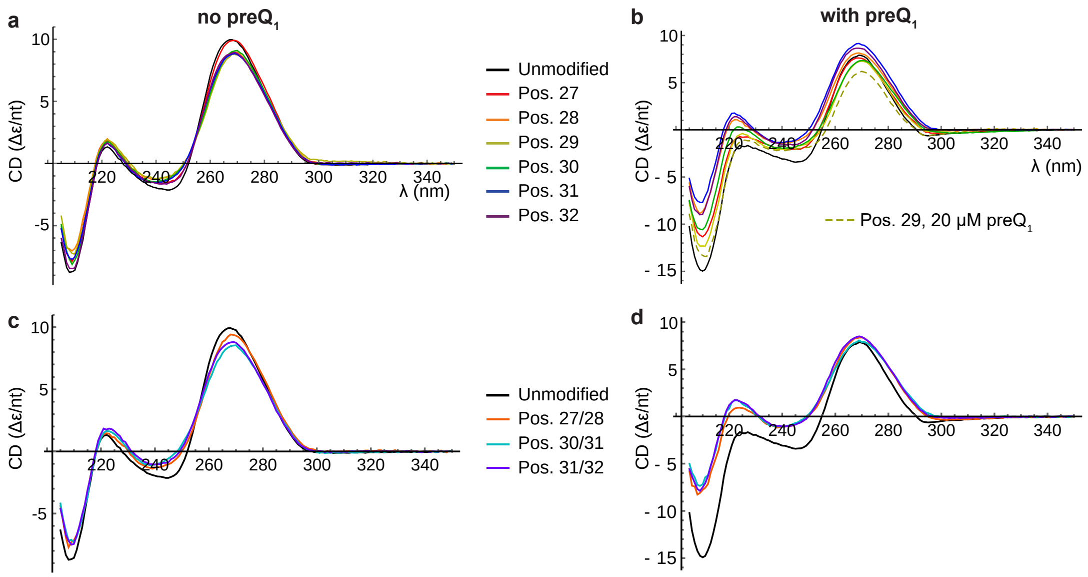


**Figure S6.** (a-b) CD spectra of unmodified and singly-labeled variants in the absence (a) or presence (b) of 5 µM preQ$_1$. Panel b additionally shows the spectrum of the position 29 variant in the presence of 20 µM preQ$_1$. (c-d): CD spectra of unmodified and doubly-labeled variants in the absence (c) or presence (d) of 5 µM preQ$_1$. These spectra were used to construct the difference spectra shown in Fig. 3 and Fig. S9.

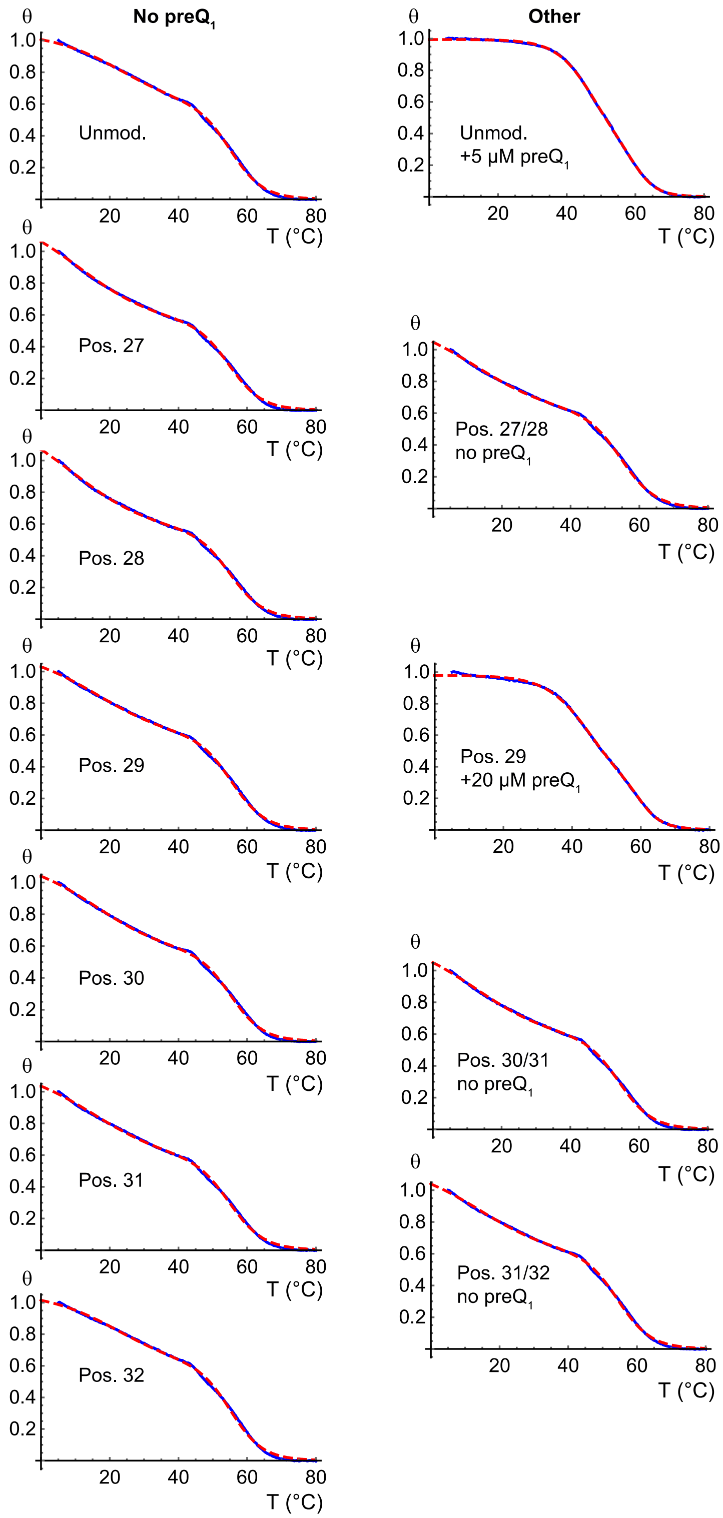


**Figure S7.** Processed melting curves (blue) and fits (dashed red) for one replicate plotted as θ (see methods) vs. T. The left column contains curves recorded on unmodified and singly-labeled riboswitch variants in the absence of $preQ_1$, and the right column contains other variants and $preQ_1$ concentrations as indicated on the plots.

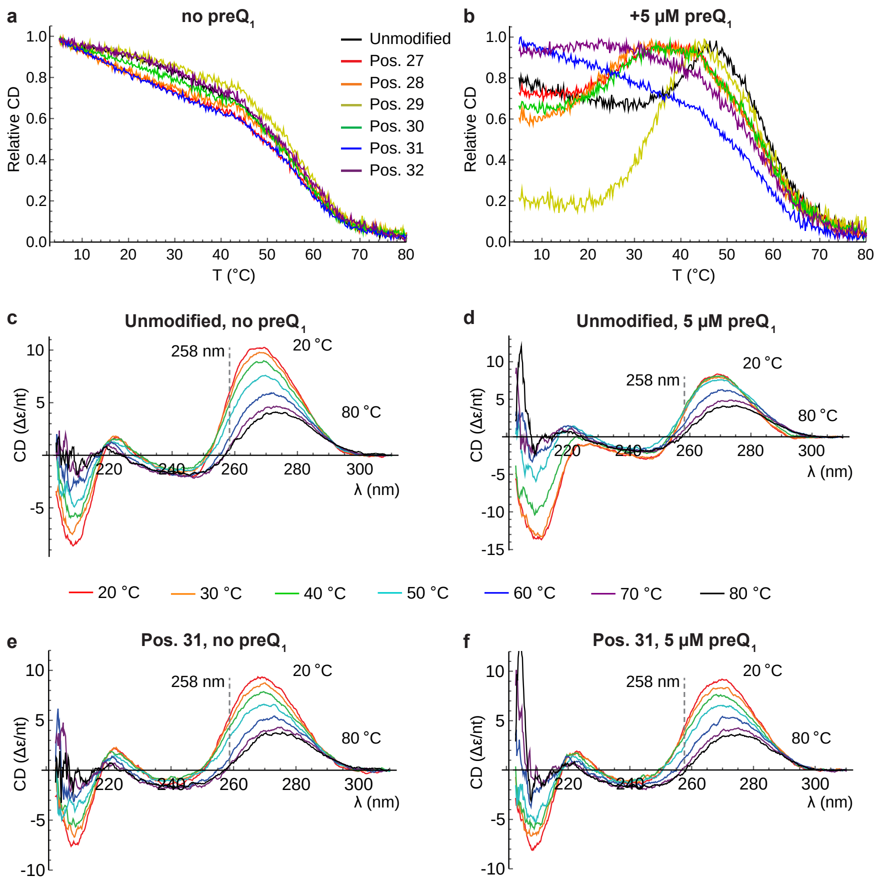


**Figure S8.** CD melting curves of unmodified and singly-labeled variants recorded at 258 nm in the absence of $Mg^{2+}$ and the absence (a) or presence (b) of 5 µM $preQ_1$. For ease of comparison, the curves were shifted and normalized to run from a minimum of 0 to a maximum of 1. (c-f) CD spectra recorded during thermal ramping at 20 (red), 30 (orange), 40 (green), 50 (cyan), 60 (blue), 70 (purple) and 80 (black) °C. The wavelength at which the CD was monitored while recording melting curves is indicated by a dashed line. (c) Unmodified riboswitch in the absence of $preQ_1$. (d) Unmodified riboswitch in the presence of 5 µM $preQ_1$. (e) Position 31 variant in the absence of $preQ_1$. (f) Position 31 variant in the presence of 5 µM $preQ_1$.

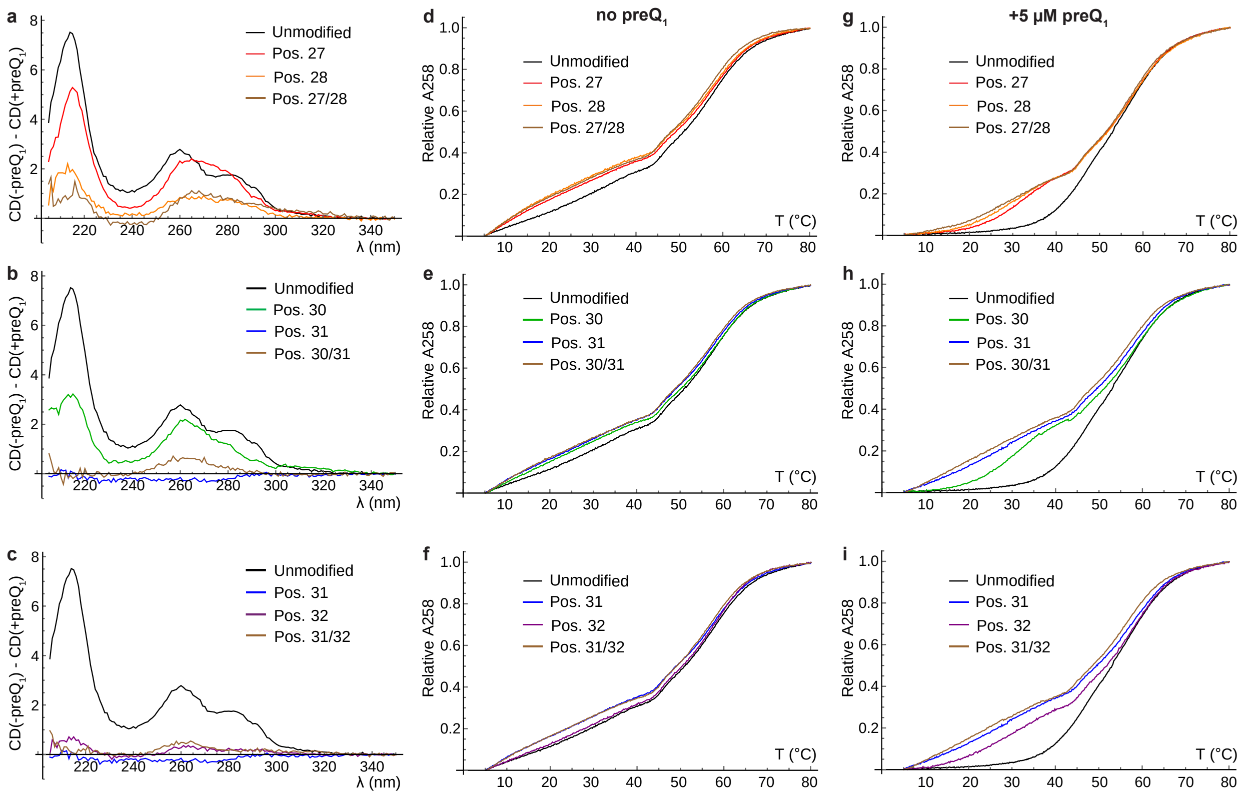


**Figure S9.** CD and thermal denaturation analysis of doubly-labeled variants. (a) CD difference spectra for unmodified, position 27, position 28, and position 27/28 variants. (b) Corresponding plots for positions 30 and 31. (c) Corresponding plots for positions 31 and 32. (d-f) Melting curves for unmodified, singly-labeled and doubly-labeled variants in the absence of $preQ_1$ (d) Positions 27 and 28. (e) Positions 30 and 31. (f) Positions 31 and 32. (g-i) Corresponding melting curves in the presence of 5 µM $preQ_1$. For ease of comparison, the melting curves were shifted and normalized to run from a minimum of 0 to a maximum of 1.
